# Exact and WKB-approximate distributions in a gene expression model with feedback in burst frequency, burst size, and protein stability

**DOI:** 10.1101/2020.10.27.357368

**Authors:** Pavol Bokes

## Abstract

The expression of individual genes into functional protein molecules is a noisy dynamical process. Here we model the protein concentration as a jump–drift process which combines discrete stochastic production bursts (jumps) with continuous deterministic decay (drift). We allow the drift rate, the jump rate, and the jump size to depend on the protein level to implement feedback in protein stability, burst frequency, and burst size. We specifically focus on positive feedback in burst size, while allowing for arbitrary autoregulation in burst frequency and protein stability. Two versions of feedback in burst size are thereby considered: in the first, newly produced molecules instantly participate in feedback, even within the same burst; in the second, within-burst regulation does not occur due to the so-called infinitesimal delay. Without infinitesimal delay, the model is explicitly solvable; with its inclusion, an exact distribution to the model is unavailable, but we are able to construct a WKB approximation that applies in the asymptotic regime of small but frequent bursts. Comparing the asymptotic behaviour of the two model versions, we report that they yield the same WKB quasi-potential but a different exponential prefactor. We illustrate the difference on the case of a bimodal protein distribution sustained by a sigmoid feedback in burst size: we show that the omission of the infinitesimal delay overestimates the weight of the upper mode of the protein distribution. The analytic results are supported by kinetic Monte-Carlo simulations.

## 1 Introduction

Molecular noise in gene expression is transmitted downstream to the level of the protein product [1, 2]. Consequently, the amount of protein exhibits broad temporal fluctuations in single cells and varies across populations of isogenic cells [3, 4]. Mathematical modelling quantifies the effects of molecular dynamics on the protein level distributions. Popular modelling frameworks include discrete-state Markov processes [5–8] and hybrid models such as piecewise-deterministic Markov processes [9–12].

Bursty protein production is a dominant source of stochastic noise in the expression of individual genes [13, 14]. Following previous studies, we model the temporal dynamics of a bursty protein by a continuous-time continuous-state drift–jump process with stochastic jumps modelling production and deterministic drift accounting for protein decay [15]. The Chapman–Kolmogorov equation for the drift–jump model with exponentially distributed burst sizes and feedback in burst frequency was formulated and solved in [16]. An exact simulation algorithm for the model and its multi-species extensions was implemented in [17]. The adjoint equation was formulated and first passage times were calculated in [18]. Numerical methods for solving the forward equation were implemented in [19, 20]. Convergence towards the steady-state solution was established by entropy methods in [21].

The drift–jump model has been extended in a number of ways, e.g. with decoy binding sites [22, 23], multiple gene copy numbers [24], time-dependent decay and production rates [25], and explicit cell partitioning [26]. The availability of explicit stationary distributions for drift-jump models facilitates calculating the Fisher information [27] and solving noise-minimisation problems [28].

Although most analyses focus on feedback in burst frequency, there are well documented examples of proteins which regulate their own stability or burst size [29–31]. Noise-reduction performance of negative feedback of the different kinds was probed using the drift–jump modelling framework in [32, 33]. Feedback in burst size manifests in the jump kernel of the drift–jump process. Different kernels have been formulated based on different physical assumptions in [34]: the “infinitesimally delayed” kernel applies if the size of a burst is regulated only by protein molecules which have been present before the burst began; the “undelayed” kernel applies if one allows regulation on the (infinitesimal) intra-burst timescale. Elongation and maturation delays make it unlikely that a protein would be effective immediately after the transcription initiation event; for this reason, the “infinitesimally delayed” kernel is deemed more realistic in [34]. The noise-reduction performance of feedback in burst size deteriorates if infinitesimal delay is accounted for [34].

The aforementioned studies extend the original explicit stationary distribution result to a generalised problem which includes not only feedback in burst frequency but also in protein stability and also undelayed feedback in burst size. However, explicit distribution is unavailable with infinitesimally delayed feedback in burst size, except for a special case of Michaelis–Menten-type response [35]. In the absence of explicit solutions, asymptotic approximations come to the fore [36, 37]. The Wentzel–Kramers–Brillouin method provides uniform approximations to stationary distributions of stochastic models that operate near the deterministic limit [38]. In drift–jump processes, deterministic behaviour is reached in the limit of very frequent and small jumps [39].

The main result of this paper is the WKB approximate distribution for a drift–jump gene expression model with positive feedback in burst size and general autoregulation in burst frequency and protein stability. The model is formulated, and the difficulties introduced by regulation of burst size are explained, in Section 2. Infinitesimal delay is identified as the source of these difficulties in Section 3. Explicit steady-state distribution is found for a modified model from which the infinitesimal delay is extricated in Section 4. Deterministic limit is identified, and its relationship with the steady-state distribution is characterised, in Section 5. The main result, the WKB approximation to the steady-state distribution of the original model, is reached in Section 6. The usefulness of the WKB solution is demonstrated on an example of a bistable switch in Section 7. The paper is wrapped up in Section 8.

## 2 Model formulation

Protein dynamics is given by the dynamic balance of production and decay. In our chosen framework, production is modelled by jumps and decay by drift of a Markov drift–jump process (Figure 1, lower left). The forward evolution of probability distributions of a Markov process is described by a Chapman– Kolmogorov equation. For drift–jump Markov processes, this takes the form of an integro-differential equation, here

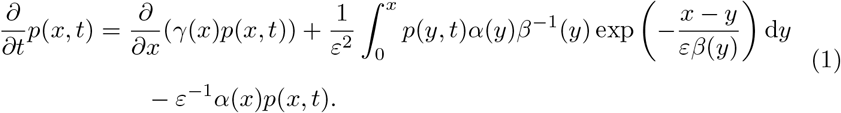

**Figure 1:**
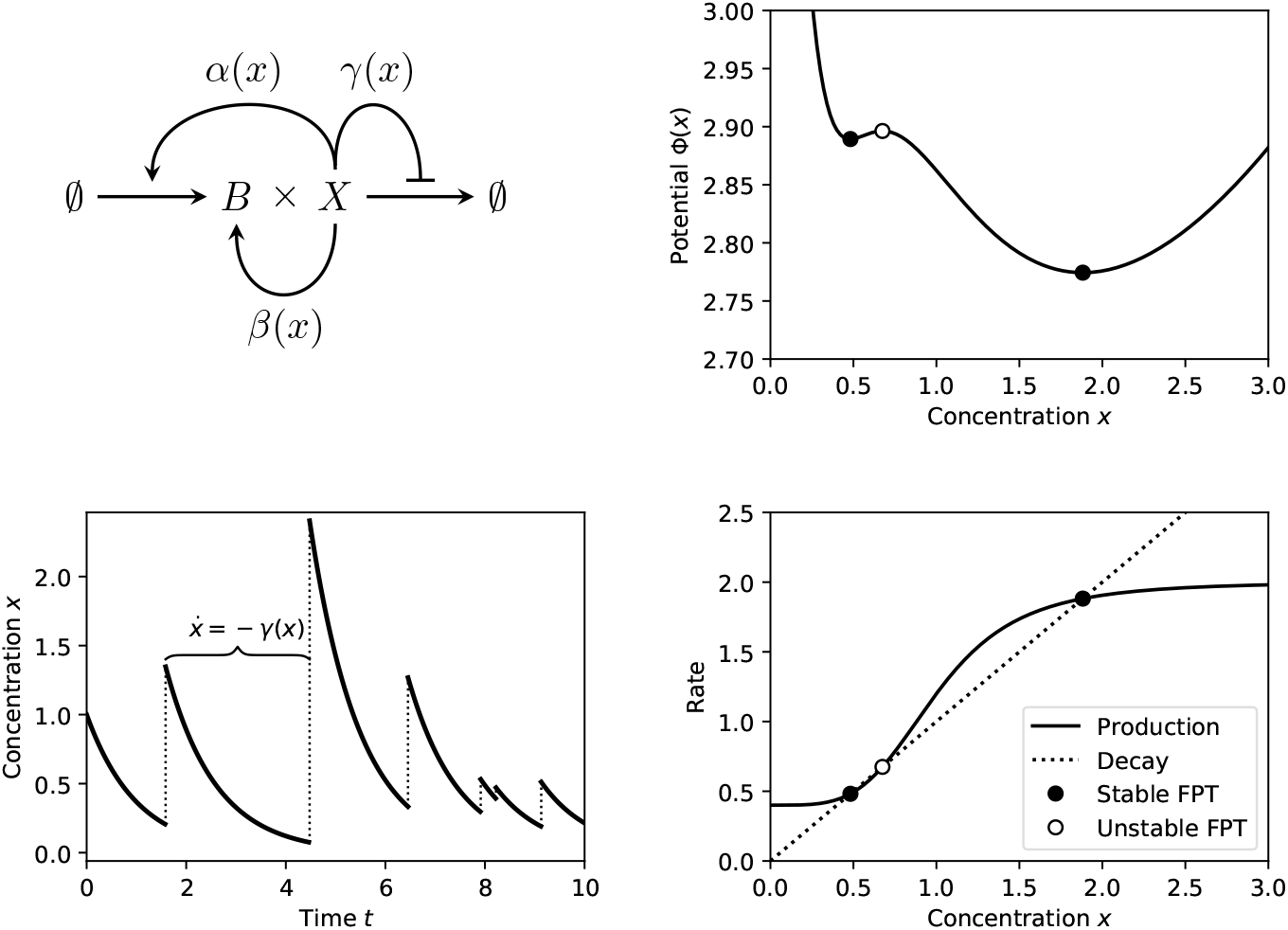
*Upper Left:* The model includes bursty protein production and continuous protein decay, and allows for feedback in burst frequency, burst size, and protein stability, as quantified by functions *α*(*x*), *β*(*x*), and *γ*(*x*), respectively. *Bottom Left:* Bursts lead to an instantaneous increases in protein concentration; between bursts protein concentration decays continuously. *Bottom Right:* In the deterministic limit *ε →* 0, the concentration changes per unit time by the difference of the production rate *α*(*x*)*β*(*x*) (solid line) and the decay rate *γ*(*x*) (dotted line). The intersections of the two are the fixed points (FPTs) of the deterministic model. *Upper Right:* The distribution potential (16) is a Lyapunov function of the deterministic model: it possesses minima/maxima where the deterministic model exhibits stable/unstable points. The depicted example pertains to feedback in burst size, with *γ*(*x*) = *x, α*(*x*) *≡* 1, *β*(*x*) = 0.4 + 1.6*x*^4^*/*(1 + *x*^4^).

The drift term accounts for the transfer of probability by deterministic decay. The integral term accounts for non-local probability transport from *y* into *x* by instantaneous production bursts; the exponential distribution of burst sizes manifests in the integral kernel. The negative term is due to transport away from *x* by bursts. Burst frequency *α*(*x*), the mean burst size *β*(*x*), and degradation rate *γ*(*x*), can depend on *x* in an arbitrary fashion (with reasonable restrictions to guarantee existence, uniqueness, and ergodicity of solutions [21]). They can be chosen so as to implement feedback loops (Figure 1, upper left). In the absence of any regulation, when *α*(*x*) and *β*(*x*) are constant and *γ*(*x*) is linear, solving (1) at steady state gives the gamma probability distribution [16]. The parameter *ε* is a scaling parameter, the decreasing of which makes bursts smaller and more frequent. In the absence of regulation, doing this leads to the Gaussian limit of the gamma distribution [34].

Following [32], we rewrite (1) as a probability conservation equation

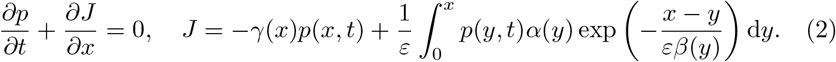

If 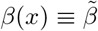, the dependence of the integral kernel 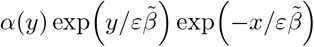 on *x* and *y* is separated, which enables to reduce the steady-state Volterra integral equation *J* = 0 into an ordinary differential equation of the first order [40]. An alternative derivation of the stationary distribution uses the Laplace transform and is based on 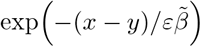 being a translational kernel [16]. If *β*(*x*) is non-constant, *x* and *y* cannot be separated in the integral kernel, nor is it a translational kernel, and an explicit solution is, to the author’s best knowledge, in general unavailable.

## 3 Removing infinitesimal delay

In this section we attribute the non-separability of the integral kernel in (2) to infinitesimal delay and formulate the undelayed model with a separable kernel. Consider time *t* of a production burst, and denote by *X*(*t*^*−*^) the state of the protein process right before and by *X*(*t*^+^) the state of the process right after the burst. Equation

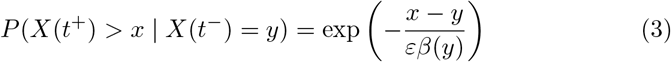

relates these random variables to the burst kernel in (2). Equation (3) is equivalent to

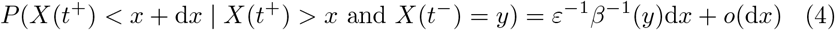

for the probability that a burst, having passed protein level *x*, will be aborted before reaching *x* + d*x*. In probability/reliability theory, this is referred to as the hazard function [41]. Importantly, the hazard function *ε*^*−*1^*β*^*−*1^(*y*) in (4) is constant with respect to the current protein level *x* and depends only on the pre-burst level *y*. This means that newly produced proteins are not engaged in feedback within the burst in which they were produced, an effect which we refer to as the infinitesimal delay.

The infinitesimal delay is eliminated from (4) by making the hazard function depend on the current protein level rather than pre-burst level, i.e.

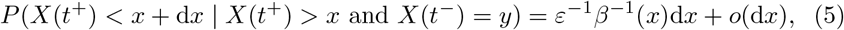

which leads to a modified burst kernel

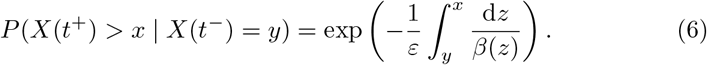

In contrast to the original kernel (3), the modified kernel (6) is a product kernel (see the next section for an explicit solution). Other notable differences are that the burst size distribution is not exponential in (6) and the function *β*(*x*) does not give the mean burst size in the modified kernel (6) like it did in the original exponential kernel (3). The only possible interpretation of *β*(*x*) in (6) is the reciprocal of the hazard rate for burst abortion.

## 4 Explicit solution to the undelayed model

We modify the master equation (2) by replacing the infinitesimally delayed burst kernel (3) by the undelayed one (6), obtaining

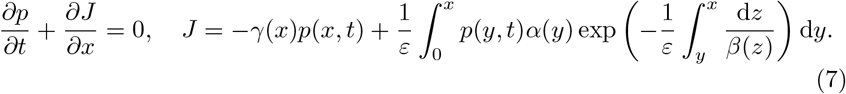

The stationary solution *p*(*x, t*) = *p*(*x*) to (7) satisfies *J* = 0, i.e. an integral equation

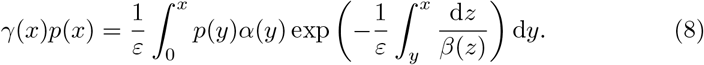

Denoting

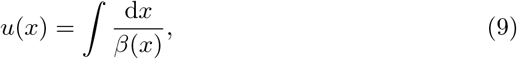

we can rewrite (8) into

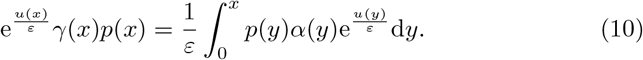

Differentiating (10) with respect to *x* yields

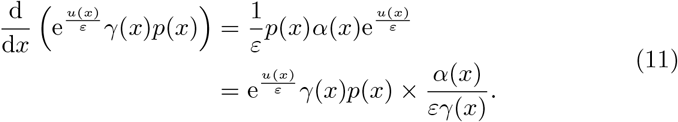

Solving the linear first-order differential equation (11) in 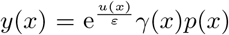 yields

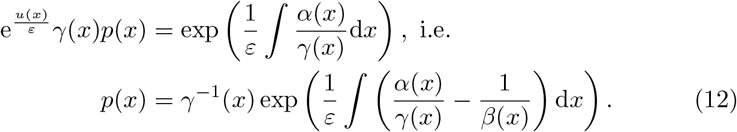

The formula (12) provides the explicit steady-state protein distribution in the modified model obtained by eliminating the infinitesimal delay in burst size.

## 5 Lyapunov function of the deterministic flow

The deterministic limit of our stochastic model (either formulation) is obtained by making the bursts extremely small and frequent, i.e. taking *ε →* 0. The nonlocal part of the probability flux *J*, whether given by (2) or (7), is dominated by contributions from a *x − y* = *O*(*ε*) boundary layer. Estimating the integral using the Laplace method [36], we find that the non-local term is at the leading order local; dropping higher-order terms yields a limiting equation

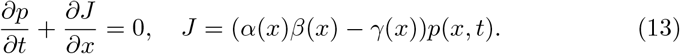

Equation (13) is a Liouville equation [39] that characterises the evolution of a distribution of particles each evolving deterministically according to

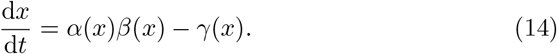

The interpretation of equation (14) is clear: in the deterministic limit, the rate of change in protein level is the burst frequency times burst size minus decay rate. Notably, both infinitesimally delayed and undelayed formulations lead to the same deterministic flow (14).

In connection with the obtained deterministic reduction, it is instructive to rewrite the explicit stationary distribution (12) as

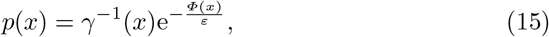

with

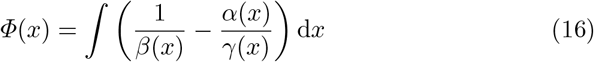

being referred to as the potential. It is clear that

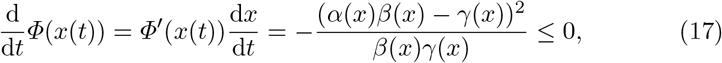

with equality in (17) holding if *x*(*t*) = *x* is a fixed point of the deterministic flow (14). Thus, the distribution potential (16) is a Lyapunov function of (14), and has local minima (maxima) where the deterministic flow (14) has stable (unstable) fixed points. The dominant contribution towards the total probability mass (15) will thereby come from the neighbourhoods of points where the potential is minimal. Therefore, the probability density function (15) is for *ε ≪* 1 sharply peaked around the stable fixed points of the deterministic flow (14).

## 6 WKB solution to the infinitesimally delayed model

Having solved the modified, undelayed master equation (8) in Section 4 and having established the relationship between the small-*ε* behaviour of the explicit solution (12) and the deterministic flow (14) in Section 5, we now proceed to analyse the stationary behaviour of the original, undelayed model. At steady state, the flux *J* in (2) vanishes, so that the stationary distribution *p*(*x*) satisfies the integral equation

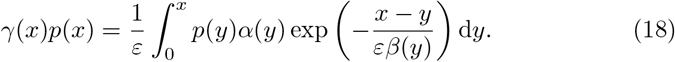

Because the integral kernel in (18) is not of product type (nor it is a translational kernel), an explicit solution is, to the author’s best knowledge, unavailable. Here we construct an *ε →* 0 asymptotic solution to (18) that applies for non-decreasing *β*(*x*), which allows for positive feedback in burst size; feedback in protein stability and burst frequency can be of either kind. Inspired by the form of the explicit solution (12) to the undelayed problem, we seek an approximate solution to (18) in the form of

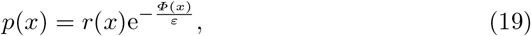

where *r*(*x*) *>* 0 and *Φ*(*x*) are functions to be determined (however, we reveal in advance that the ensuing calculations will demonstrate that *Φ*(*x*) coincides with the potential (16) identified in the explicit solution (12)). The ansatz (19) is referred to in literature as the WKB asymptotic form after Wentzel, Kramers, and Brillouin [42]; we thereby assume a regular dependence

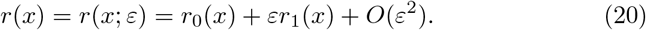

Without any loss of generality, *Φ*(*x*) is assumed to be independent of *ε*. In the context of WKB asymptotics, *Φ*(*x*) is referred to as the WKB quasi-potenial, which we will shorten to potential, and *r*(*x*) as the exponential prefactor, or simply the prefactor.

Inserting (19) into (18) yields

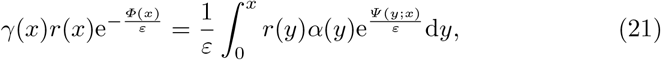

where

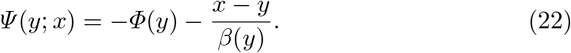

Estimating the integral on the right-hand side of (21) by means of per-partes integration yields (the dash represents the derivative w.r.t. *y* throughout)

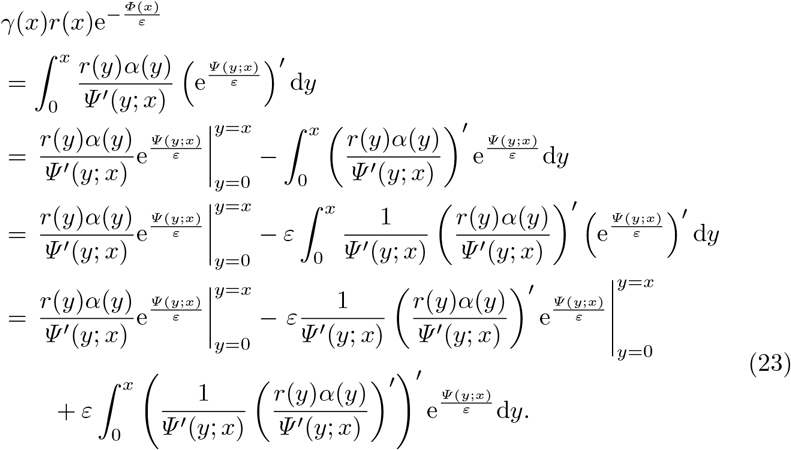

The contributions to (23) from the *y* = *x* boundary dominate the contributions away from the boundary provided that *Ψ ′* (*y*; *x*) *>* 0: the assumption will be verified post hoc provided that *β*(*x*) is non-decreasing^1^. Further iteration of per-partes integration shows that the integral term in (23) is 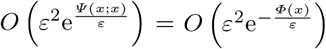 [36]. Neglecting the higher-order terms and cancelling 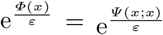 on either side of the equation, we find

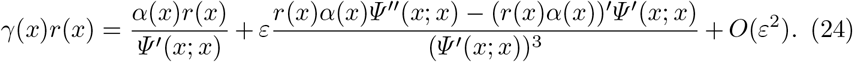

Inserting (20) into (24) and collecting terms of same order yields

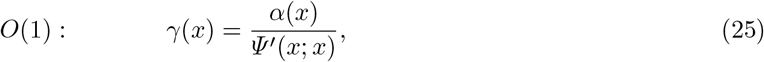

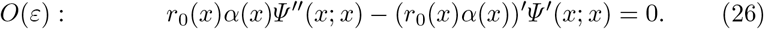

Differentiating (22) with respect to *y* gives

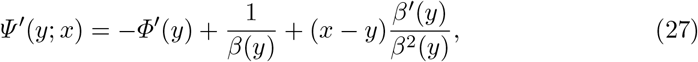

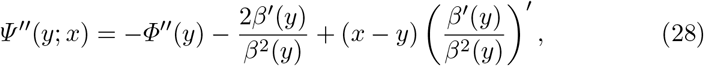

from which at *y* = *x* we obtain

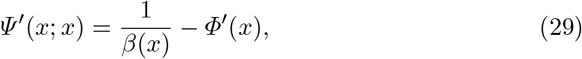

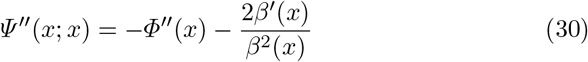

for the coefficients in (25)–(26). Combining (29) and (25) yields

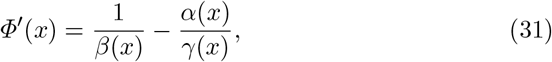

which determines (up to an additive constant) the potential in the WKB solution (19). Inserting (31) into (27) verifies that, provided that *β*(*x*) is non-decreasing, then *Ψ* (*y*; *x*) is an increasing function of *y*, which was assumed in the approximations (23). Importantly, (31) is the same potential as identified in the exact solution to undelayed model, cf. (15)–(16).

In order to determine, to the leading order, the prefactor of the WKB solution, we insert (31) into (30) to find that

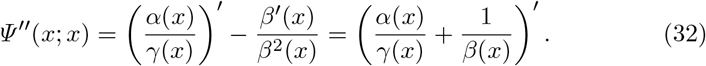

Simplifying (26) using (25) and (32) yields

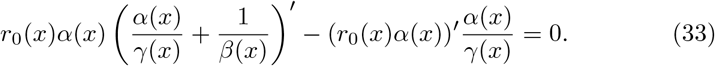

Solving the linear first-order differential equation (33) in the unknown *y*(*x*) = *r*_0_(*x*)*α*(*x*) gives the prefactor in the form

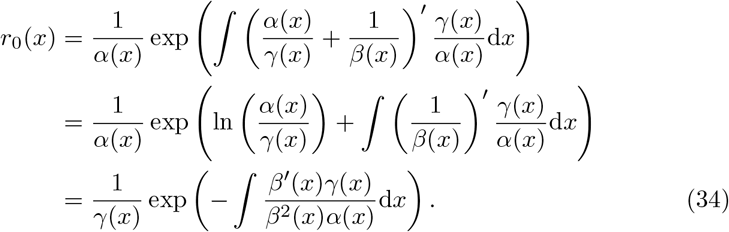

Contrastingly, the prefactor in the explicit solution to the undelayed model is simply given by 1*/γ*(*x*), cf. (15). Therefore, provided that *β*(*x*) *≠* const, i.e. there is feedback in burst size, the infinitesimally delayed and undelayed model differ in the exponential prefactor.

## 7 Bistable switch

In this section we provide an example in which the WKB stationary distribution

- accurately describes the large-time behaviour of the (infinitesimally delayed) model;
- is substantially different from the exact stationary distribution (based on disregarding the infinitesimal delay).

By constructing such an example, we will demonstrate that there is a a nonempty subset of the parametric space, in which the new result (the WKB distribution) applies whereas the previously available result (the exact distribution of the undelayed model) fails to do so.

Preliminary considerations narrow down the space in which such an example can be found; first, as the WKB result is an approximation, the perturbation parameter *ε* has to be small; second, the limiting deterministic equation has to have multiple fixed points. To see that, assume to the contrary that the deterministic model has a single globally stable equilibrium *x*_*∗*_. This point also represents the unique global minimum *x*_*∗*_ of its Lyapunov function, i.e. the potential *Φ*(*x*). Performing a parabolic approximation around *x*_*∗*_ in (15) or (19) shows that the distribution is approximately normally distributed with mean *x*_*∗*_ and variance *ε/Φ″* (*x*_*∗*_) [44] (this procedure is equivalent with the linear noise approximation [45, 46]). At the leading order, the distribution thus depends only on the potential (which is the same whether the infinitesimal delay is included or omitted) but not on the prefactor (which differentiates the two results). Therefore, there will not be a substantial difference between the WKB distribution and the exact distribution of the undelayed model if the deterministic dynamics possesses just one steady state.

Consider, on the other hand, a bistable switch with two stable fixed points *x*_*±*_ (separated by an unstable one). The parabolic approximation argument implies that the distribution will be close to a mixture of two Gaussians with means *x*_*±*_ and variances *ε/Φ″* (*x*_*±*_). Importantly, the mixture weights will depend not only on the potential but also on the prefactor [47]. Therefore, the two model versions (infinitesimally delayed and undelayed) are expected to give different weights to the two modes, and result into substantially different distributions.

By Section 6, in order that the prefactors of the two distributions be different, there has to be feedback in burst size, but burst frequency and protein stability can be unregulated. In the absence of regulation, burst frequency is constant and the decay rate is proportional to the protein concentration, and without loss of generality we set

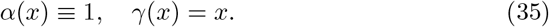

With (35), bursts arrive as per a Poisson process with rate *ε*^*−*1^, and the sample paths are proportional to e^*−t*^ on any time interval between two consecutive bursts.

In order to sustain bistability, regulation in burst size has to be sufficiently sigmoid, such as given by the widely-used Hill function

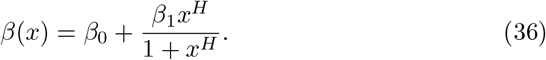

With (35)–(36), the deterministic model (14) is given by

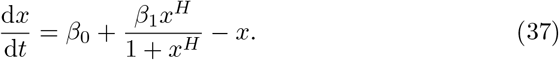

If *H >* 1, there exist *β*_0_ *>* 0 and *β*_1_ *>* 0 such that (37) has two stable fixed points separated by an unstable one (cf. Figure 1, lower right).

We validate our theoretical results by histograms of multiple independent runs of an exact simulation algorithm of the protein dynamics. In the absence of regulation in burst frequency and protein stability, the simulation algorithm is remarkably simple, and works in the following way. Assume that the sample path *x*(*t*) has already been generated on an interval 0 *≤ t ≤ t*_cur_ (initially *t*_cur_ = 0 and *x*(0) = *x*_0_ is an initial value). The iteration of the algorithm requires to draw two random variates, *θ* and 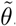, from the uniform distribution in the unit interval. The first is used to generate, by the inversion sampling method, an exponentially distributed waiting time *τ* = *−ε*ln*θ* until next burst; the sample path is set to 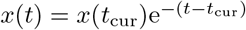 for *t*_cur_ *< t < t*_cur_ + *τ* (the time of the next burst excepted). The second random variate is used to generate the exponentially distributed burst size 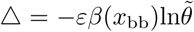, where *x*_bb_ = *x*(*t*_cur_)e^*−τ*^ is the concentration immediately before the burst; at the time of the next burst the sample path is set to *x*(*t*_cur_ + *τ*) = *x*_bb_ + Δ. The iteration of the algorithm is concluded by updating the current time according to *t*_cur_ *← t*_cur_ + *τ*. The algorithm is iterated until the path *x*(*t*) is sampled on a time interval (0, *t*_end_) (in practice on a discretisation of the interval). The algorithm is exact in the sense that it does not introduce any truncation error; the only sources of error are statistical and round-off errors.

Figure 2, upper panels, show sample paths on a moderate timescale (left panel) and on an extremely slow timescale (right panel). The stochastically generated sample paths are compared to the deterministic solutions to (37) (initiated to the same values *x*_0_). The fixed points of (37) are also shown as horizontal lines. On the moderate timescale, the stochastic process fluctuates around the deterministic solutions; on the large timescale, the process transitions, with a very small rate, between the basins of attractions of the two stable fixed points.

**Figure 2:**
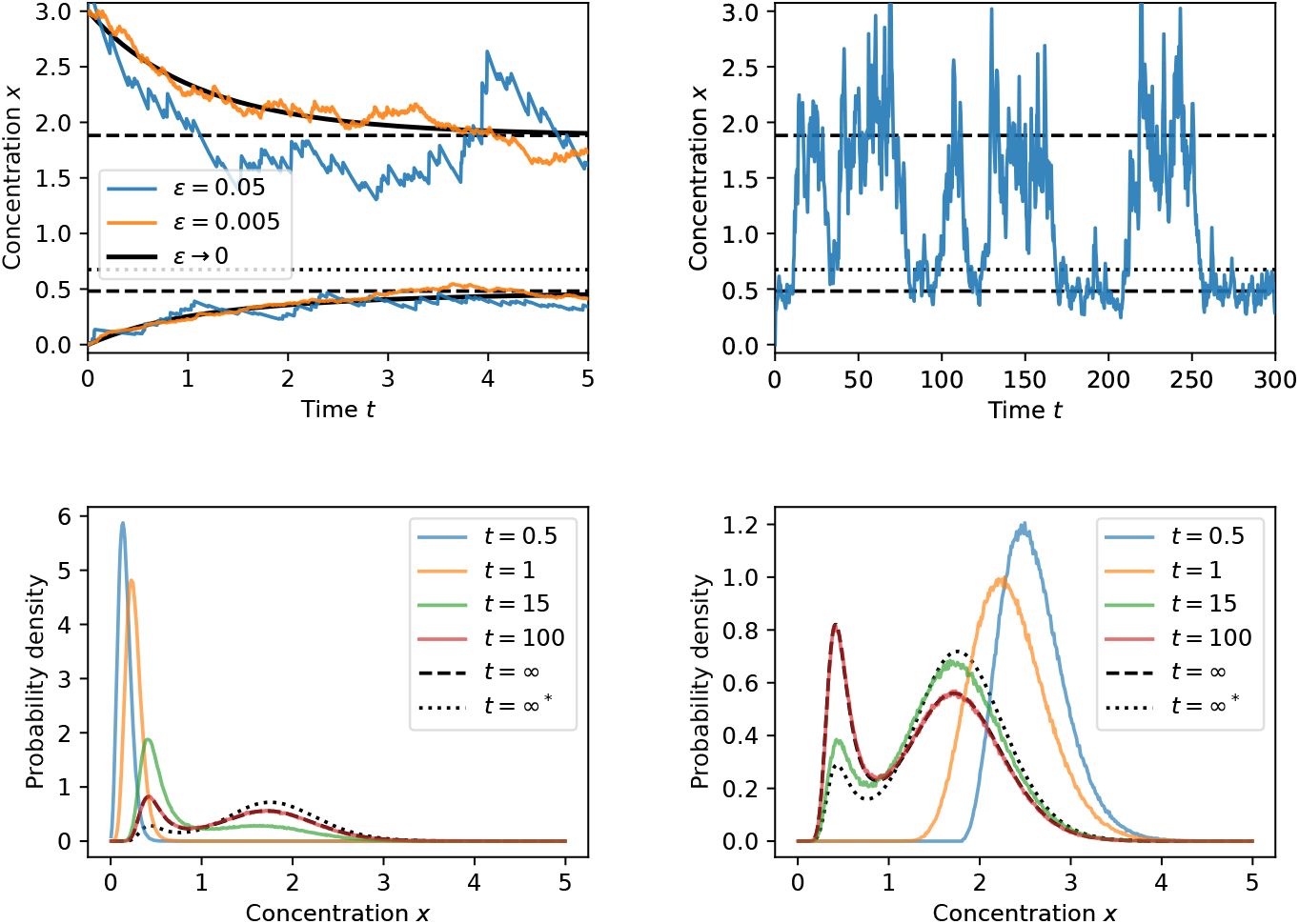
*Upper Left:* For small values of the noise parameter *ε*, the sample paths of the stochastic model (coloured curves) are close to the solutions of the deterministic model (black curves). The stable/unstable fixed points of the deterministic model (14) are shown as dashed/dotted horizontal lines. *Upper Right:* Transitions between the basins of attractions of the stable steady states occur on an extremely slow timescale. *Lower Panels:* Simulation-based time-dependent distributions (coloured curves) approach, as simulation time increases, the WKB stationary distribution (solid black curve). This is markedly different from the exact stationary distribution of the model without infinitesimal delay (dashed black curve). The initial condition is *x*_0_ = 0 (left panel) and *x*_0_ = 3 (right panel). *Parameter values:* Feedback is in burst size, with *α*(*x*) *≡* 1, *β*(*x*) = 0.4 + 1.6*x*^4^*/*(1 + *x*^4^), *γ*(*x*) = *x*. The noise parameter is fixed to *ε* = 0.05 in all panels except the upper left, where it is varied (the selected values are given inset).

Figure 2, lower panels, compare simulation-based probability densities to the theoretical results for two initial conditions *x*_0_ = 0 (left panel) or *x*_0_ = 3 (right panel). The densities were estimated from *N* = 10^6^ independent sample paths *x*_1_(*t*), …, *x*_*N*_ (*t*) evaluated at selected time values *t ∈ {*0.5, 1.0, 15, 100*}*. For each of these time points, a histogram was calculated by dividing the concentration interval (0, 5) into 500 equally sized bins and counting the number of realisations in each bin. For the final time point, *t* = 100, the empirical histogram (red colour) is in excellent agreement with the WKB stationary distribution (black solid line). The latter is given by (19) (the WKB form), (16) (the potential), and (34) (the prefactor), and was evaluated by numerical quadrature. Contrastingly, the exact stationary distribution (dashed black line), as given by (15)–(16), of the undelayed model underestimates the lower mode and overestimates the upper mode of the distribution.

## 8 Discussion

This paper considered a drift–jump model for gene expression that includes feedback in burst frequency, burst size, and protein stability. Two versions have been formulated that differ in the inclusion/omission of infinitesimal delay in burst size. The undelayed model admits an explicit solution [cf. 32, 33]. In the near-deterministic asymptotic regime of small and frequent bursts, the explicit solution assumes a WKB-type dependence on the perturbation parameter. For the infinitesimally delayed model, an exact solution is unavailable, but we constructed a WKB approximation in the near-deterministic regime. We thereby focused on the case of positive feedback in burst size (but allowing for an arbitrary regulation of burst frequency and protein stability). We show that the solutions of the two model versions have the same WKB quasi-potential but differ in the exponential prefactor.

That the two models share the same potential is not entirely surprising given that they generate the same deterministic flow in the limit of very frequent and small bursts. Nevertheless, although sharing the same deterministic flow implies that both potentials, both acting as Lyapunov functions, have their extrema located at the same points, the equality of potentials is not a-priori clear. Indeed, there are well documented examples, notably a discrete-state Markov chain and its Kramers–Moyal diffusion truncation, which reduce to the same deterministic flow, but generate markedly different quasi-potentials [48]. Therefore, the equality of potentials implies that the closeness of the two models goes beyond sharing a common deterministic limit.

Comparison of the analytic results to stochastic simulations showed that the WKB solution provides reliable approximations to the large-time distribution of protein concentration. Contrastingly, the exact distribution of the undelayed model can give different weights to the peaks of a bimodal protein distributions. Hence, we demonstrated the usefulness and applicability of WKB methods in a hybrid, drift–jump, modelling framework. The study also clarified the origins and consequences of infinitesimal delay in feedback in burst size.

## Footnotes

1 The case of a decreasing *β*(*x*) is dealt with in a follow-up manuscript [43].

